# BEACON: Benchmark for Comprehensive RNA Tasks and Language Models

**DOI:** 10.1101/2024.06.22.600190

**Authors:** Yuchen Ren, Zhiyuan Chen, Lifeng Qiao, Hongtai Jing, Yuchen Cai, Sheng Xu, Peng Ye, Xinzhu Ma, Siqi Sun, Hongliang Yan, Dong Yuan, Wanli Ouyang, Xihui Liu

## Abstract

RNA plays a pivotal role in translating genetic instructions into functional outcomes, underscoring its importance in biological processes and disease mechanisms. Despite the emergence of numerous deep learning approaches for RNA, particularly universal RNA language models, there remains a significant lack of standardized benchmarks to assess the effectiveness of these methods. In this study, we introduce the first comprehensive RNA benchmark BEACON (**BE**nchm**A**rk for **CO**mprehensive R**N**A Task and Language Models). First, BEACON comprises 13 distinct tasks derived from extensive previous work covering structural analysis, functional studies, and engineering applications, enabling a comprehensive assessment of the performance of methods on various RNA understanding tasks. Second, we examine a range of models, including traditional approaches like CNNs, as well as advanced RNA foundation models based on language models, offering valuable insights into the task-specific performances of these models. Third, we investigate the vital RNA language model components from the tokenizer and positional encoding aspects. Notably, our findings emphasize the superiority of single nucleotide tokenization and the effectiveness of Attention with Linear Biases (ALiBi) over traditional positional encoding methods. Based on these insights, a simple yet strong baseline called BEACON-B is proposed, which can achieve outstanding performance with limited data and computational resources. The datasets and source code of our benchmark are available at https://github.com/terry-r123/RNABenchmark.

## 1 Introduction

RNA plays a vital role in numerous biological processes, including protein synthesis, enzymatic activities, and gene regulations [16, 47, 73, 56]. Unlike its more famous counterpart DNA, RNA is not restricted to information storage but actively participates in translating genetic instructions into functional proteins, modulating gene expression through various mechanisms, and regulating cellular responses to internal and external stimuli [27]. As a dynamic intermediary between DNA and protein, RNA governs crucial biological processes, making it a focal point of research in molecular biology and biomedicine. Consequently, understanding the diverse functions of RNA is crucial to unraveling the complexities of cellular processes and deciphering the underlying mechanisms of diseases.

Despite its critical importance, understanding the functional roles of RNA poses significant challenges. Inspired by the success of machine learning in various fields, there have been extensive research efforts in recent years to apply machine learning approaches to RNA tasks. Initially, traditional machine learning algorithms such as support vector machine and random forest paved the way for predictive modeling in RNA studies [38, 66, 41]. The evolution of deep learning, especially through Convolutional Neural Networks (CNNs), has enabled more nuanced analyses of RNA sequences and structures [59, 6, 35]. More recently, pre-trained language models (LM) have revolutionized RNA research, facilitating more accurate predictions of RNA function and interactions [8, 12, 76]. These advancements significantly deepen our understanding of RNA’s regulatory roles in cellular processes.

According to the central dogma of molecular biology [14], genetic information flows unidirectionally from DNA to RNA and then to protein, or directly from RNA to protein. While established benchmarks for DNA [29, 44] and protein [51, 75] have significantly aided research in these areas, RNA, a crucial component of the central dogma, lacks such standardized benchmarks. Therefore existing RNA models are often evaluated using disparate individual datasets, making it difficult to conduct fair comparisons between different methods and hindering the development of the field.

To address this gap, we present the first comprehensive RNA benchmark called BEACON (**BE**nchm**A**rk for **CO**mprehensive R**N**A Task and Language Models). As shown in Fig. 1, BEACON contains a curated collection of 13 important RNA-related tasks derived from a comprehensive review of RNA-related research papers [18, 35, 13], containing 967k sequences with lengths ranging from 23 to 1182. These tasks cover sequence-level and nucleotide-level analyses across three main fields: **Structural Analysis** focuses on deciphering RNA’s secondary structures and three-dimensional configurations, essential for its interactions with other molecules and for therapeutic design. **Functional Studies** investigate RNA’s roles in gene regulation and its implications for disease, which are vital for protein translation and treatment of disease. **Engineering Applications** explore RNA’s potential in synthetic biology to enhance its utility in biotechnology and medicine, exploring how RNA can be utilized to solve complex biological challenges.

**Figure 1.**
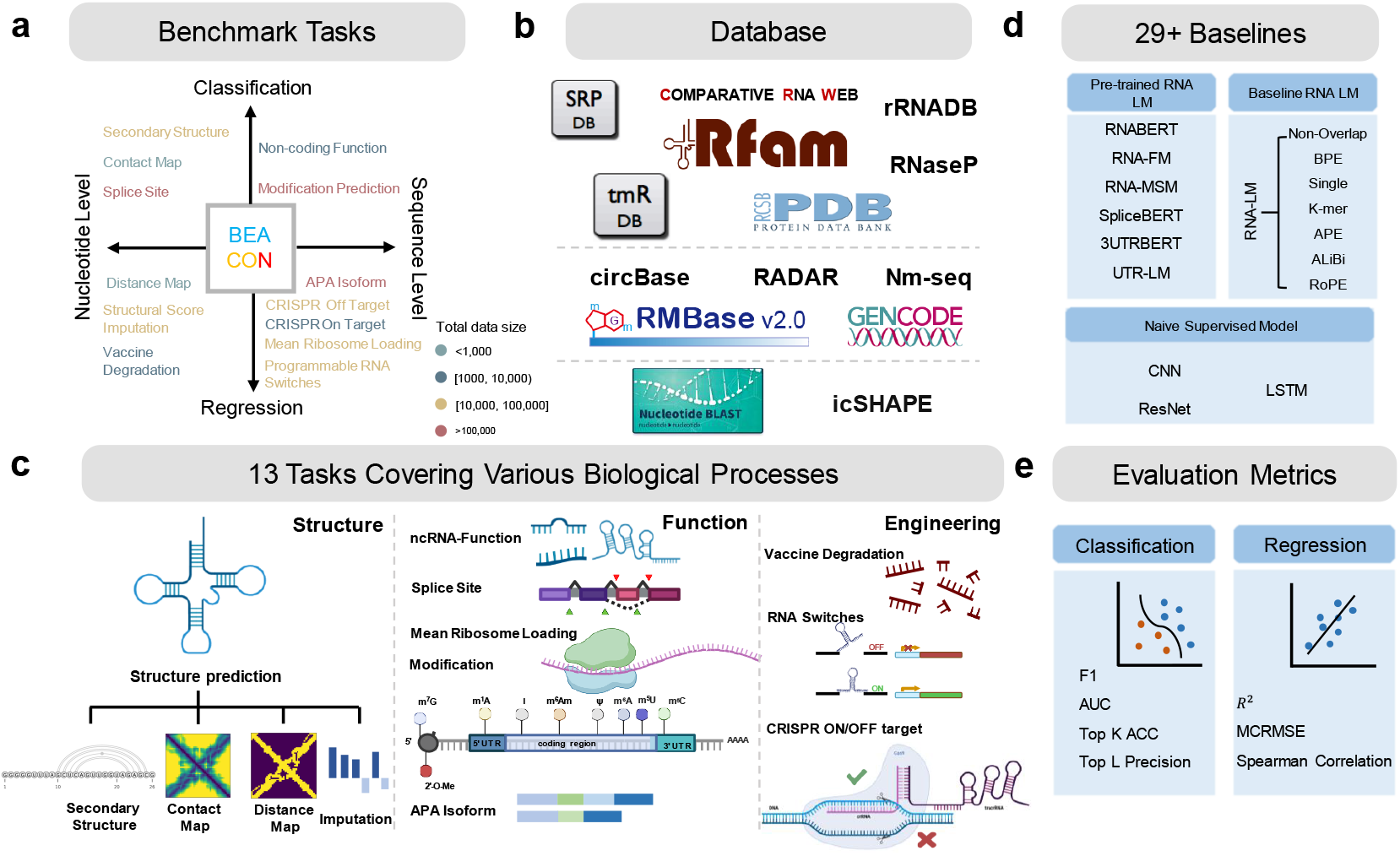
Overview of BEACON: **a:** Categorization of the 13 benchmark tasks into classification and regression at both nucleotide and sequence levels. **b:** Diverse database distinguished by data size and source type. **c:** Visual representations of tasks across Structure, Function, and Engineering. **d:** List of baseline models, including naive supervised deep models and advanced RNA language models. **e:** Metrics for evaluating model performance in classification and regression tasks, tailored to RNA analysis specifics.

In addition, we evaluate a diverse range of models using our benchmark, including traditional models like CNNs, ResNets, and LSTMs, as well as advanced RNA foundation models like RNA-FM [8] and UTR-LM [12]. Surprisingly, ResNet and LSTM are proved to be strong baselines, managing to outperform language models on several tasks. Additionally, pretrained RNA language models surpassed previous task-specific state-of-the-art (SOTA) performances on 8 out of the 13 tasks, demonstrating significant potential. Next, we explore the impact of various components in RNA language models for the community, with a particular focus on tokenization methods and positional encodings. We conclude some experimental findings, based on which we further propose a robust yet efficient baseline, BEACON-B, that incorporates Attention with Linear Biases (ALiBi) and single nucleotide tokenization, providing an extremely fast and easy-to-use open-source pre-training model for the community.

Overall, our contributions can be summarized as follows:

- We establish the first comprehensive benchmark for RNA research with 13 diverse tasks, covering structure, function, and engineering aspects.
- We conduct a thorough evaluation of pre-trained RNA language models, providing insights into their strengths and limitations across different tasks.
- We investigate the component impacts of RNA language models in depth, and propose BEACON-B, a simple yet strong baseline, that benefits subsequent research in the field.

## 2 Related Works

### RNA tasks

RNA research is categorized into three primary areas: structure, function, and engineering. Structural tasks, such as predicting secondary structures [18] and contact maps [67], aim to understand RNA configurations. Functional tasks focus on the biological roles of RNA, including splice site prediction [35] and non-coding RNA function classification [2]. Engineering tasks involve designing RNA molecules with specific properties for applications in synthetic biology, such as discriminating programmable RNA switches [4].

### Deep learning methods in RNA tasks

Deep learning has been pivotal in addressing these tasks. For structural predictions, U-Net [54] has been employed to model secondary structures [25]. In functional studies, methods like SpliceAI utilize dilated convolutions for effective splice detection [35]. For engineering challenges, LSTMs have been used to design programmable RNA switches, demonstrating their versatility in handling complex sequence data [4]. The development of foundational RNA models like RNA-FM [8], RNA-BERT [1], RNA-MSM [78], SpliceBERT [10], UTR-LM [12] and 3UTRBERT [76] represents a significant advancement, capable of tackling multiple RNA-related tasks by leveraging advanced language modeling techniques. These models promise a broader understanding of RNA biology. However, they often lack thorough evaluations across different tasks, highlighting a gap in the systematic assessment of their capabilities. For instance, UTR-LM [12] only focuses on 5’ UTR function-related tasks. This limitation underscores the need for robust, cross-disciplinary evaluation approaches to fully explore and utilize the potential of these models in RNA research.

### Benchmarks in molecular biology

While AI for RNA research is a relatively new field and lacks comprehensive benchmarks, numerous benchmarks have been established for DNA [29, 17, 45] and protein [51, 75, 48, 26] studies. Grešováet al. proposed Genomic Benchmarks [29], which includes a collection of genomic sequence classification tasks. Marin et al. constructed BEND [45], a comprehensive benchmark for DNA, encompassing tasks such as gene finding, enhancer annotation, histone modification, CpG methylation, etc. Notin et al. introduced ProteinGym [48], a benchmark specifically designed for protein fitness prediction and design, while Gao et al. proposed ProteinBench [26], focusing on protein design. Additionally, Xu et al. developed PEER [75], a comprehensive protein benchmark involving function, localization, structure, etc. As AI for RNA research develops, some preliminary benchmarks in RNA have emerged. RnaBench [57] is the benchmark for computational RNA modeling, though it only involves tasks related to RNA secondary structure and RNA design. Many real-life applications, such as RNA-based therapeutics, require a comprehensive understanding of the functions of ncRNA and mRNA [80]. To address this gap, we developed BEACON, a comprehensive benchmark for RNA that covers a wide range of tasks covering structural analysis, functional studies, and engineering applications.

## 3 Benchmark Tasks

BEACON comprises 13 tasks designed to evaluate RNA models comprehensively, covering structural analysis, functional studies, and engineering applications. The following sections provide detailed information, including data statistics, evaluation metrics, and data sources as shown in Tab 1.

**Table 1:** Overview of the 13 benchmark tasks across three major RNA task groups. Nucleotide- level tasks require labels to have the same length as input sequences. Sequence-level tasks require nucleotides from one input sequence to share one label. Cls and Reg denote classification and regression, respectively. MCRMSE means Mean Columnwise Root Mean Squared Error.

| RNA Task | #Train/Validation/Test | Metric | Task Type | RNA Level | Max/Mean Length | Source/Venue |
| --- | --- | --- | --- | --- | --- | --- |
| **Structure** |  |  |  |  |  |  |
| SSP | 10,814/1,300/1,305 | F1 | Multi-label Cls | Nucleotide | 499/133.8 | bpRNA/NAR [18] |
| CMP | 188/23/80 | Top L Precision | Multi-label Cls | Nucleotide | 960/110.3 | RNAcontact/BIOINF [39] |
| DMP | 188/23/80 | $R^2$ | Reg | Nucleotide | 960/110.3 | RNAcontact/BIOINF [39] |
| SSI | 14,049/1,756/3,095 | $R^2$ | Reg | Nucleotide | 100/100 | StructureImpute/NMI [28] |
| **Function** |  |  |  |  |  |  |
| SPL | 144,628/18,078/16,505 | Top-k ACC | Multi-class Cls | Nucleotide | 100/100 | SpliceAI/Cell [35] |
| APA | 145,463/33,170/49,755 | $R^2$ | Reg | Sequence | 186/186 | APARENT/Cell [6] |
| ncRNA | 5,679/650/2,400 | ACC | Multi-class Cls | Sequence | 1182/158.4 | Noorul's/NMI [2] |
| Modif | 304,661/3,599/1,200 | AUC | Multi-label Cls | Sequence | 101/101 | MultiRM/NC [63] |
| MRL | 76,319/7,600/7,600 | $R^2$ | Reg | Sequence | 100/61.5 | Optimus/NBT [59] |
| **Engineering** |  |  |  |  |  |  |
| VDP | 2,155/245/629 | MCRMSE | Multi-label Reg | Nucleotide | 130/118.5 | OpenVaccine/NMI [71] |
| PRS | 73,227/9,153/9,154 | $R^2$ | Multi-label Reg | Sequence | 148/148 | Angenent-Mari's/NC [3] |
| CRI-On | 1,453/207/416 | Spearman Corr | Reg | Sequence | 23/23 | DeepCRISPR/GB [13] |
| CRI-Off | 14,223/2,032/4,064 | Spearman Corr | Reg | Sequence | 23/23 | DeepCRISPR/GB [13] |

Each molecule is assigned a categorical label *y* ∈ {0, 1, …, 12} to denote its function. The dataset [2, 24] comprises contributions from GENCODE, circBase, and Rfam, encompassing various ncRNAs. Accuracy (ACC) at the sequence level is the evaluation metric.

### 3.1 Structure Prediction

**Secondary Structure Prediction (SSP)** identifies paired regions (stems) and unpaired regions (loops, bulges, junctions) within RNA molecules. The target matrix is *y* ∈ ℝ^*l*×*l*^ indicating whether each nucleotide pair forms a base pair as part of the RNA’s secondary structure. We adopt the bpRNA-1m database [18], which contains detailed annotations of over 100,000 single-molecule RNA structures. The evaluation metric is the F1 score.

*Impact:* Accurate secondary structure prediction is pivotal for elucidating the structural and functional dynamics of RNA. By precisely mapping these structures, researchers gain insights into functional regions and interaction sites, contributing significantly to areas such as drug discovery and genetic research.

**Contact Map Prediction (CMP)** identifies pairs of nucleotides in RNA that are in close proximity in their three-dimensional structures. Each nucleotide pair is associated with a binary label *y* ∈ {0, 1} indicating whether they contact (within a distance threshold of 8 Å). Following [67], we utilize a dataset derived from non-redundant RNA 3D structures documented by [39], and evaluate predictions using the Top-L precision metric.

*Impact:* Accurately identifying nucleotide interactions is pivotal for inferring the tertiary structure of RNA molecules. These predictions enhance our understanding of RNA folding and function, contributing to advancements in RNA-based therapeutics and biotechnology.

**Distance Map Prediction (DMP)** estimates the physical distances between pairs of nucleotides within an RNA molecule. The target distance matrix *y* ∈ ℝ^*l*×*l*^ records the distance between every pair of nucleotides within the sequence. The same dataset as the contact map prediction task is used, with *R*^2^ serving as the evaluation metric.

*Impact:* Inter-nucleotide distance prediction offers detailed spatial information, facilitating the construction of accurate three-dimensional models by providing distance restraints.

**Structural Score Imputation (SSI)** predicts missing structural information within RNA molecules, with each nucleotide assigned with an experimentally derived structural score *y* ∈ *R*. The dataset [28] is derived from icSHAPE sequencing data of the HEK293 cell line, with 30% of nucleotides randomly masked as null in the training set, and downsampling leading to missing values in 3,095 fragments in the testing set. The evaluation metric is *R*^2^.

*Impact:* Accurate structural score imputation provides enhanced and comprehensive structural information, crucial for the development of RNA-based therapeutics and diagnostics. Improved structural data enable precise targeting of RNA molecules in disease treatment, potentially leading to more effective interventions.

### 3.2 Function Prediction

**Splice Site Prediction (SPL)** classifies each base within a sequence into one of three categories: acceptor (a), donor (d), or neither (n), with the categorical label *y* ∈ {0, 1, 2}. The task uses Jaganathan’s dataset [35] with Top-k accuracy as the evaluation metric.

*Impact:* Splice site prediction is crucial for studying gene expression and regulation within biological systems. Accurate prediction of splice sites helps determine the precise locations within the genome where splicing occurs, enabling the detection of non-coding genomic variations that could impact protein synthesis, particularly those resulting in cryptic splicing.

**APA Isoform Prediction (APA)** predicts the usage ratio of the proximal polyadenylation site (PAS) in the 3’ untranslated region (3’ UTR) for each variant, recorded in target *y* ∈ ℝ. We filter 228k sequences from over 3 million APA reporter gene data from Bogard’s dataset [6], this regression task assesses the proportion of proximal APA isoforms. The evaluation metric is the *R*^2^ value.

*Impact:* APA is a common gene expression regulation mechanism that generates different RNA transcripts and protein isoforms by modulating RNA 3’ UTR processing [77]. This regulatory method can affect gene expression levels and functions, playing a crucial role in cellular biological processes and development.

**Non-coding RNA Function Classification (ncRNA)** classifies ncRNA molecules into categories like microRNAs (miRNAs), long non-coding RNAs (lncRNAs), and small interfering RNAs (siRNAs).

*Impact:* Classifying ncRNA functions is crucial for understanding their diverse roles in gene regulation and cellular processes. Accurate classification enhances our knowledge of regulatory networks and aids in elucidating disease mechanisms. This contributes to identifying new biomarkers and therapeutic targets, advancing molecular biology research, and improving disease diagnosis and treatment.

**Modification Prediction (Modif)** predicts twelve widely occurring types of RNA modifications from a given RNA sequence, indicated by a categorical label *y* ∈ {0, 1, …, 11}. We adopt Song’s dataset that contains 20 epi-transcriptome profiles for 12 different types of RNA modifications obtained from 15 base-resolution technologies, where over 300,000 sites were collected and divided into training, validation, and test sets. We use AUC as the metric.

*Impact:* Post-transcriptional RNA modifications enhance the structural and functional diversity of RNA molecules, impacting all stages of RNA life [22]. Due to the complex and diverse characteristics of RNA sequences, different modifications may correspond to distinct sequence features. Accurately identifying RNA modification sites is crucial for understanding the functions and regulatory mechanisms of various RNAs.

**Mean Ribosome Loading (MRL)** predicts the MRL value for a given sequence, with target *y* ∈ ℝ representing the level of mRNA translation activity into proteins. Data from Reid’s dataset [59] of 91,519 5’ UTR sequences and their variants are used to calculate the MRL for each sequence. The model’s performance is evaluated using the *R*^2^ value.

*Impact:* MRL refers to the average ribosome load on a specific mRNA sequence under given conditions, indicating the translation efficiency of ribosomes on that mRNA. Modulating the features and structures of the 5’ UTR sequence can influence ribosome loading on mRNA, thereby regulating protein expression levels [40, 7].

### 3.3 Engineering Prediction

**Vaccine Degradation Prediction (VDP)** forecasts the stability and shelf life of vaccines under different environmental conditions. For each nucleotide, the three properties are recorded in target *y* ∈ ℝ^3^. We use data from the “Stanford OpenVaccine” [71] competition on Kaggle and the RNA design platform Eterna, which includes detailed measurements for 6,043 diverse RNA constructs. The evaluation metric is the Mean Columnwise Root Mean Squared Error (MCRMSE).

*Impact:* Accurate predictions of vaccine degradation under different environmental conditions are crucial for optimizing storage and transportation protocols, ensuring vaccines remain potent until administration. Enhanced degradation predictions are particularly beneficial for distributing vaccines in challenging environments, such as resource-limited settings, by providing guidelines to maintain vaccine stability and efficacy.

**Programmable RNA Switches (PRS)** involves identifying synthetic RNA molecules that can alter their conformation and function in response to specific signals. The target *y* ∈ ℝ ^3^ records the ON, OFF and ON/OFF states activity given an RNA sequence. The dataset, analyzed by Angenent-Mari [3], includes 91,534 toehold switches in vivo, covering 23 viral genomes and 906 human transcription factors, with GFP signal intensity measurements indicating ON and OFF states activity levels [4]. The *R*^2^ metric evaluates the effectiveness of these switches.

*Impact:* Programmable RNA switches provide precise control of gene expression and cellular functions, serving as powerful tools for investigating biological processes [64, 46]. In therapeutic applications, these switches hold promise for developing targeted and personalized treatments by responding to disease-specific signals, offering innovative approaches to medical intervention.

**CRISPR On-Target Prediction (CRI-On)** evaluates the efficiency of single-guide RNAs (sgRNAs) directed by Cas proteins in gene editing within specific target sites. Each sgRNA’s knockout efficacy is quantified and presented as target *y* ∈ ℝ. The dataset [13] comprises approximately 15,000 sgRNAs targeting 1,071 genes across four different cell lines, with performance evaluated using the Weighted Spearman correlation coefficient.

*Impact:* CRISPR-Cas technology has transformed genetic engineering with significant enhancements in genome editing accuracy and safety. Effective on-target predictions are essential for designing sgRNAs that precisely modify genetic sequences without affecting unintended regions, thus improving therapeutic outcomes and research accuracy [74, 33].

**CRISPR Off-Target Prediction (CRI-Off)** assesses the likelihood and frequency of CRISPR-induced mutations at unintended genomic locations. The efficacy of sgRNA specificity is quantified using a target *y* ∈ ℝ, capturing the frequency of off-target cleavage. The evaluation dataset [13] contains data for about 160,000 potential off-target sites across 30 sgRNAs in various cell types, with the Weighted Spearman correlation coefficient serving as the metric.

*Impact:* Precision in off-target predictions is critical for advancing CRISPR technology by reducing unintended genetic modifications, which can lead to harmful effects. Accurate off-target analysis helps refine sgRNA designs, enhancing the safety and efficacy of CRISPR applications in clinical settings and research.

## 4 Models

We consider three types of baseline models in our benchmark, including naive supervised models, pre-trained language models, and the proposed BEACON-B. We give the details in the following part and summarize them in Tab. 2.

**Table 2:** Detailed specifications and pre-training data of RNA language models analyzed in the study.

| RNA Foundation Model | Number of Parameters (M) | Max Token length | Pre-trained Data | Tokenizer | Positional Encoding |
| --- | --- | --- | --- | --- | --- |
| RNA-FM [8] | 99.52 | 1024 | Multispecies ncRNA [68] | Single | APE |
| RNA-BERT [1] | 0.48 | 440 | Human ncRNA [68] | Single | APE |
| RNA-MSM [78] | 95.92 | 1024 | Homologous sequences [37, 9] | Single | APE |
| SpliceBERT-H510 [10] | 19.45 | 510 | Human pre-mRNA [30] | Single | APE |
| SpliceBERT-MS510 [10] | 19.45 | 510 | Multispecies pre-mRNA [30] | Single | APE |
| SpliceBERT-MS1024 [10] | 19.72 | 1024 | Multispecies pre-mRNA [30] | Single | APE |
| UTR-LM-MRL [12] | 1.21 | 1026 | Multispecies 5'UTR [15, 58, 7] | Single | RoPE |
| UTR-LM-TE&EL [12] | 1.21 | 1026 | Multispecies 5'UTR [15, 58, 7] | Single | RoPE |
| 3UTRBERT-3mer [76] | 86.14 | 512 | Human 3'UTR [32] | K-mer | APE |
| 3UTRBERT-4mer [76] | 86.53 | 512 | Human 3'UTR [32] | K-mer | APE |
| 3UTRBERT-5mer [76] | 88.45 | 512 | Human 3'UTR [32] | K-mer | APE |
| 3UTRBERT-6mer [76] | 98.05 | 512 | Human 3'UTR [32] | K-mer | APE |

### Naive Supervised Models

We utilize three widely-used sequence encoders: CNN [62], ResNet [51], and LSTM [51]. We mainly follow the design choices described in [75], employing 2 layers for CNN, 8 resblocks for ResNet, and 3 Bi-LSTM layers for LSTM, with 5.4M, 11M, and 26.7M parameters, respectively.

### Pre-trained Language Models

We evaluate the performance of several language models, including RNA-FM [8], RNABERT [1], RNA-MSM [78], SpliceBERT [10], 3UTRBERT [76], and UTRLM [12]. These models vary significantly in size, ranging from 0.48M to 99.52M parameters, and are pre-trained on diverse RNA data sources including ncRNA, pre-mRNA, mRNA-3’UTR, and mRNA-5’UTR. For consistency, we choose to fine-tune them using identical settings.

### Baseline RNA LM and BEACON-B

We conduct ablation studies on two key aspects of RNA LM: 1) tokenization methods including Single Nucleotide (Single), Byte-Pair Encodings (BPE) [61, 79], Overlapping K-mer (K-mer, we use 6mer for experiments) [76, 36] and Non-overlapping K-mer (Non-overlap) [17] 2) positional encodings including Absolute Positional Encodings (APE) [21], Attention with Linear Biases (ALiBi) [50] and Rotary Positional Encodings (RoPE) [65]. The findings indicate that single nucleotide tokenization outperforms both K-mer, BPE, and Non-overlap, and ALiBi shows advantages over both RoPE and APE. Consequently, we propose a robust yet efficient BEACON baseline (BEACON-B) that incorporates single nucleotide tokenization and ALiBi as positional encodings, based on the BERT backbone.

## 5 Results

### 5.1 Training setups

To ensure a fair comparison, we fully fine-tune all the BERT-like RNA foundation models including RNA-FM, RNABERT, RNA-MSM, SpliceBERT, 3UTRBERT, UTR-LM and BEACON-B under the same training settings. For simple supervised methods (CNN, ResNet and LSTM) and baseline RNA LM, we train them from scratch using similar training settings. For each model, we search for its learning rate from 1e-5 to 5e-3. All experiments are repeated with three random seeds, and we report the average performance alongside sample standard deviations. More details are in Appendix A.1.

### 5.2 Task Pipeline

Our approach incorporates three pipelines for different types of tasks in the BEACON. In nucleotide-level tasks, due to the complexity of outputs in structural tasks, we further categorize the tasks of Secondary Structure, Contact Map, and Distance Map into a more detailed nucleotide-nucleotide level prediction.

**Sequence Level Prediction** For sequence-level tasks, we apply an attentive weighted sum of all nucleotides for naive supervised models and use the [CLS] token from language models. Both representations are processed through an MLP layer to derive the sequence-level predictions.

**Nucleotide Level Prediction** For tasks requiring resolution at the nucleotide level, individual representations for each nucleotide are processed through a Multilayer Perceptron (MLP) to generate nucleotide-level predictions. Specifically, the representation for a nucleotide are calculated by averaging the representations of all tokens that cover it, as illustrated in Appendix Fig. 2.

**Nucleotide-Nucleotide Relation Prediction** To analyze relationships between nucleotides, we compute a self outer product of the nucleotide representations to form a matrix that cotains the pairwise interactions between nucleotides. This matrix is then passed through a simple Resnet to get the final output.

### 5.3 Benchmark results

In Tab 3, we report the benchmark results for popular and opensource methods, including literature SOTAs, naive supervised models and existing RNA language models.

**Table 3:**
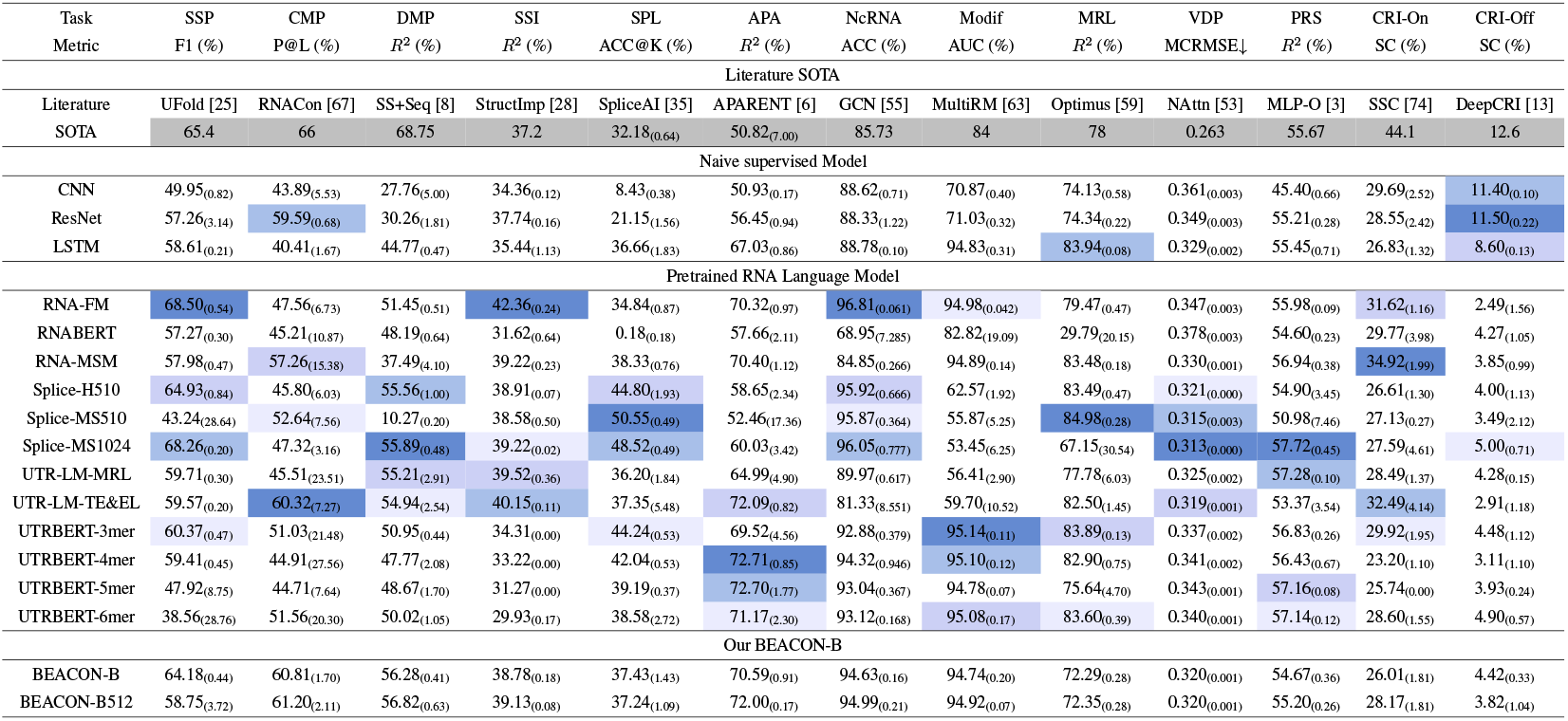
Benchmark results across various 13 RNA tasks. We use four color scales of blue to denote the first, second, third and fourth best performance among naive supervised models and pre-trained RNA LMs. Mean (std) is reported for each experiment.

#### ResNet and LSTM are strong naive supervised models

ResNet, which has only been trained on downstream tasks, can outperform most if not all language models on some tasks. LSTM outperforms the other Naive supervised Models on 9 out of 13 tasks, and the performance is better by a large margin on many tasks.

#### Pre-trained RNA language models have good potential for RNA understanding

It outperforms the previous task-specific SOTA on 8 out of 13 tasks, demonstrating that the additional unsupervised pre-training brings a lot of gains. However, there is still a long way to go on individual tasks, such as contact map prediction and distance map prediction in the structural task, vaccine degradation rate prediction in the engineering task, and CRISPR on- and off-target prediction. Of course, the previous SOTA method used additional features such as secondary structure, but it shows that there is still a lot of room for improvement in the RNA language model.

#### SpliceBERT and RNA-FM are superior models for various tasks

Both SpliceBERT-MS1024 and RNA-FM got first place in 3 out of 13 tasks, and had top performances in other tasks as well, showing they have learned rich patterns and evolution knowledge from multi-species RNA sequences.

#### Pre-training of specific RNA attributes will result in a gain on tasks with corresponding attributes

First, when specific RNA attributes are included in the pre-training it brings gains to the downstream tasks corresponding to the attributes. For example, RNA-FM pre-trained with non-coding RNA achieves the best performance in non-coding RNA family prediction, SpliceBERT pre-trained on pre-mRNA learns information about potential shear mRNAs for the best shear site prediction, and 3UTRBERT uses sequences from the 3’UTR region to learn 3’UTR function worked best in the prediction of APA isoforms in the 3’UTR functional region, and similarly, the pre-trained UTR-LM in the 5’UTR region worked well in the prediction of ribosome loading in the 5’UTR association. Second, specific attributes also give gains for having other RNA attributes, for example, 3UTRBERT, although pre-trained on 3’UTR sequences, also gained on the prediction of 5’UTR function.

### 5.4 Component Analysis of RNA language models

In Tab 4, we study the language model component effect from tokenizer and positional encoding.

**Table 4:**
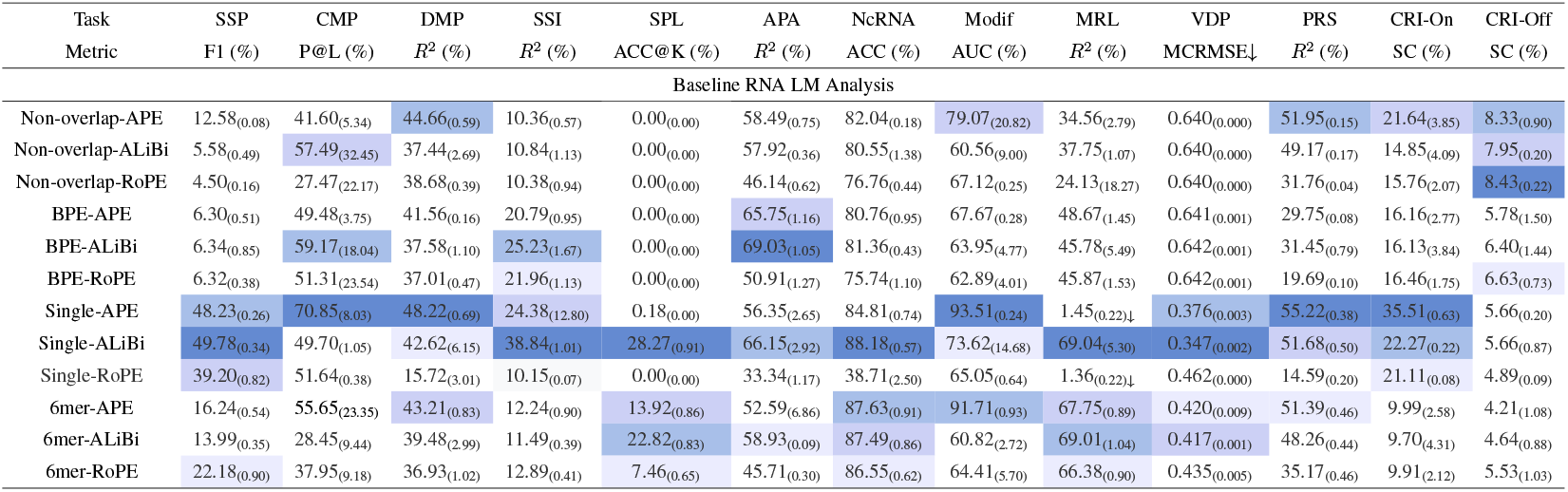
Performance of baseline RNA LMs with different tokenizers and positional encodings.

#### The single nucleotide tokenizer is a powerful tool for RNA language models

As shown in Tab 11, it achieves the best performance on 11 out of 13 tasks, significantly outperforming other tokenizers. BPE and non-overlapping tokenizers are generally ineffective at the nucleotide level, as they lose precision from overlapping. Similarly, the 6mer approach adds local information before the individual tokens, potentially introducing redundancy. We argue that the single nucleotide tokenizer can learn global information, including surrounding context, through self-attention mechanisms. Thus, using the single nucleotide tokenizer is sufficient, and future work should focus on designing interpretable tokenizers based on single nucleotide units [69, 42].

#### ALiBi is better for RNA sequences understanding

For tasks involving shorter sequences, the specific advantages of RoPE or other complex encoding schemes may not be fully realized. RoPE, which is highly effective in long sequences due to its rotational component that maintains relative positioning across long distances, might not provide significant benefits over simpler methods like ALiBi in shorter sequences. Moreover, ALiBi linearly biases the attention scores based on relative positions, which helps the model better generalize across different sequence lengths.

### 5.5 BEACON-B: an Efficient Baseline for RNA Language Models

Based on the above analysis of the different components of the RNA language model, combined with the Tab 4, we use the single nucleotide tokenizer, ALiBi as the positional encoding, and pre-train on filtered human ncRNA sequences from RNACentral [68]. We propose the low-resource and cost-effective BEACON-B (pre-trained on 1026 length seqs) and BEACON-B512 (pre-trained on 512 length seqs and FlashAttn [20]) as a baseline to provide an extremely fast and easy to use open-source pre-training model for subsequent researchers.

Although with very small GPU days (days * GPUs) as the cost of pre-training, BEACON-B can even outperform SOTA pre-trained RNA LMs on some tasks such as **contact map prediction** and **distance map prediction**. Compared with other models that also report pre-training resources, BEACON-B and BEACON-B512 can match or even surpass existing RNA language models in one-to-one comparisons on **almost half of the tasks** listed in Tab 3 and Tab 5, despite being pretrained with significantly fewer resources. This demonstrates that the insights we obtain from the important components in analysing the RNA language model are vital and that biological motifs and configurations on limited RNA data can be fully explored by utilising such a combination of components in a good way.

**Table 5:**
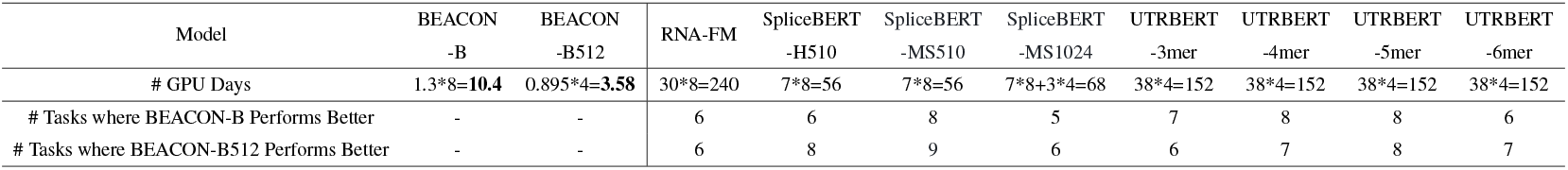
Comparison of GPU days and the number of tasks (total: 13) where BEACON-B demon- strates performance superiority over other methods.

## 6 Conclusions

### Summary

In this work, we present BEACON, the first comprehensive RNA benchmark, which encompasses 13 diverse tasks spanning structural analysis, functional studies, and engineering applications. BEACON aims to address the critical gap in standardized evaluation for RNA models. We assess various models, from traditional approaches like CNNs to advanced RNA foundation models, providing insights into their task-specific performances. Additionally, we analysis the vital components of RNA LM from tokenization and positional encoding. Building upon this, we propose BEACON-B, an efficient baseline that incorporates single nucleotide tokenization and ALiBi. BEACON’s standardized evaluation framework and the insights provided into RNA modeling components are expected to significantly advance RNA research, facilitating the development of more sophisticated models and enhancing our understanding of RNA’s diverse roles in biology.

### Limitation & future work

Despite the comprehensiveness of BEACON, it has some limitations for future work. While BEACON includes 13 diverse RNA-related tasks, it may not cover all aspects of RNA biology, necessitating the inclusion of additional tasks and datasets in future versions. The influence of pre-training datasets and hyperparameters on model performance also needs further systematic exploration to optimize configurations for specific RNA tasks. Although BEACON-B serves as an efficient baseline, there is potential for developing more advanced models that leverage RNA’s unique structural characteristics. Additionally, BEACON primarily evaluates predictive accuracy, suggesting the need to incorporate metrics like interpretability, computational efficiency, and robustness for a more holistic assessment. Addressing these limitations and exploring new directions will not only advance RNA research but also deepen our understanding of its indispensable roles in genetic regulation, disease pathogenesis, and therapeutic development.

## A Appendix / supplemental material

Optionally include supplemental material (complete proofs, additional experiments and plots) in the appendix. All such materials **SHOULD be included in the main submission**.

### A.1 Experimental settings for tasks

#### A.1.1 Most of the tasks

Most of the tasks are trained using the same training settings shown in the Tab 6. These tasks include structural score imputation, splice site prediction, APA isoform prediction, non-coding RNA function classification, modification prediction, programmable RNA switches, CRISPR on-target prediction and CRISPR off-target prediction.

**Table 6:** Configuration settings for most of the tasks training.

| config | value |
| --- | --- |
| optimizer | AdamW |
| optimizer epsilon | 1e-8 |
| optimizer momentum | $\beta_1, \beta_2 = 0.9, 0.999$ |
| weight decay | 0.01 |
| learning rate sch. | linear decay |
| learning rate | [1e-5, 5e-3] |
| warmup steps | 50 |
| epochs | 30 |
| batch size | 32 |
| gradient accumulation | 1 |
| dtype | float16 |

In particular, for the structural score imputation task, the input is the sequence accompanied by structural scores. The sequence is fed to the model and undergoes the transformation to the nucleotide-level representation. We concatenates it with the MLP-passed structural scores, and then use the regression header to get imputation scores.

For CRISPR off-target, the input is two sequences including sgRNA and target sequences. We feed them through the same model separately, then concat them together. Finally, we use the regression header to get the predicted value of off-target.

#### A.1.2 Vaccine degradation prediction task

Vaccine degradation prediction is trained using the settings shown in the Tab 7.

**Table 7:** Configuration settings for vaccine degradation prediction training.

| config | value |
| --- | --- |
| optimizer | AdamW |
| optimizer epsilon | 1e-8 |
| optimizer momentum | $\beta_1, \beta_2 = 0.9, 0.999$ |
| weight decay | 0.01 |
| learning rate sch. | linear decay |
| learning rate | [1e-5, 5e-3] |
| warmup steps | 50 |
| epochs | 100 |
| batch size | 32 |
| gradient accumulation | 1 |
| dtype | float16 |

#### A.1.3 Nucleotide-nucleotide level tasks

Nucleotide-nucleotide level tasks are trained using the settings shown in the Tab 8. In addition, the representation also follows Fig 2

**Figure 2.**
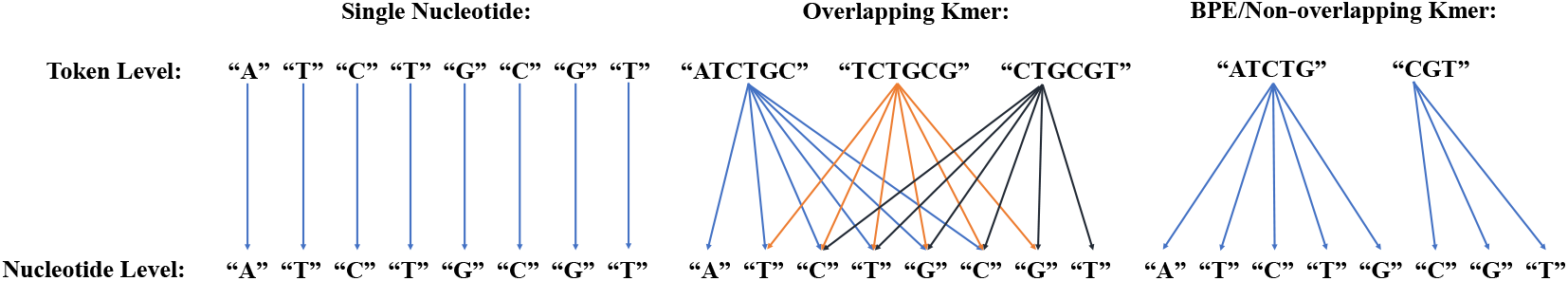
Derivation of nucleotide-level representations. In single nucleotide tokenization, a token directly corresponds to a nucleotide, thus the representations are identical. For overlapping Kmer tokenization, the nucleotide representation is the averaged representation of tokens covering it. In Byte-Pair Encoding (BPE) and non-overlapping K-mer tokenization, the representation is derived from the token covering it.

**Table 8:** Configuration settings for nucleotide-nucleotide level tasks training.

| config | value |
| --- | --- |
| optimizer | Adam |
| optimizer epsilon | 1e-8 |
| optimizer momentum | $\beta_1, \beta_2 = 0.9, 0.999$ |
| learning rate sch. | cosine decay |
| learning rate | [1e-5, 5e-3] |
| warmup epochs | 1 |
| epochs | 100 |
| batch size | 1 |
| gradient accumulation | 8 |
| dtype | float16 |

### A.2 Methods in Benchmark

All pre-trained benchmarked methods use a Masked Language Modeling (MLM) objective.

In the MLM task, a sequence is provided as input, with 15% of its tokens randomly masked. The entire masked sequence is then processed by the model, which is tasked with predicting the original tokens. This approach is analogous to the Cloze task in traditional language modeling.

- 15% of the tokens in the sequence are masked.
- In 80% of the cases, the masked tokens are replaced by a special <mask> token.
- In 10% of the cases, the masked tokens are substituted with a random token different from the original.
- In the remaining 10% of cases, the masked tokens remain unchanged.

#### A.2.1 RNABERT

##### Training Objectives

RNABERT was pre-trained with two objectives: masked language modeling (MLM) and structural alignment learning (SAL).

For SAL, the model learns to predict the structural alignment between two RNA sequences. It achieves this by being trained to predict the alignment score of RNA sequence pairs using the Needleman-Wunsch algorithm.

##### Training Data

The RNABERT model was pre-trained using a subset of 76,237 human ncRNA sequences from RNAcentral. The dataset was preprocessed by applying 10 different masking patterns to the 76,237 sequences, resulting in a final dataset comprising 762,370 sequences.

#### A.2.2 RNA-FM

##### Training Data

The RNA-FM model was pre-trained using data from RNAcentral. To ensure the dataset was non-redundant, RNA-FM applied CD-HIT (CD-HIT-EST) with a cut-off at 100% sequence identity, resulting in a final dataset containing 23.7 million unique RNA sequences.

#### A.2.3 RNA-MSM

Unlike other methods, RNA-MSM utilizes homologous sequences as input to provide additional evolutionary information, similar to MSATransformer [52].

To ensure fairness, homologous sequences were not included in the input during evaluation.

##### Training Data

RNA-MSM was pre-trained using data from Rfam, which includes homologous sequences. To prevent potential overfitting in structural inference, RNA-MSM excluded families with experimentally determined structures, such as ribosomal RNAs, transfer RNAs, and small nuclear RNAs. The final dataset comprises 3,932 RNA families, with a median of 2,184 MSA sequences per family. To augment the number of homologous sequences, RNA-MSM employed an automated pipeline, RNAcmap3 [11], for homolog search and sequence alignment.

#### A.2.4 SpliceBERT

##### Training Data

The SpliceBERT model was pre-trained using messenger RNA precursor sequences obtained from the UCSC Genome Browser.

SpliceBERT gathered reference genomes and gene annotations from the UCSC Genome Browser for 72 vertebrate species. Bedtools getfasta was used to extract pre-mRNA sequences from the reference genomes based on these gene annotations. The resulting pre-mRNA sequences were then utilized for pre-training SpliceBERT. The pre-training dataset comprises 2 million pre-mRNA sequences, with a total length of 65 billion nucleotides.

#### A.2.5 3UTRBERT

##### Training Data

The 3UTRBERT model was pre-trained using human mRNA transcript sequences obtained from GENCODE.

3UTRBERT collected 108,573 unique human mRNA transcripts from GENCODE, utilizing only the longest transcript for each gene in the pre-training process. To avoid codon constraints in the CDS region and to reduce the complexity of the full mRNA transcripts, only the 3’ untranslated regions (3’UTRs) of the mRNA transcripts were used. The average length of the 3’UTRs was 1,227 nucleotides, with a median length of 631 nucleotides. Each 3’UTR sequence was divided into non-overlapping patches of 510 nucleotides, with the remaining sequences padded to the same length.

#### A.2.6 UTR-LM

##### Training Objectives

In addition to MLM pre-training, UTR-LM employs two additional supervised objectives: Secondary Structure (SS) and Minimum Free Energy (MFE).

Both secondary structure and the MFE value are calculated using ViennaRNA [43]. To prevent information leakage, UTR-LM calculates the secondary structure loss only on the masked positions. The output embedding of the cls token is used by UTR-LM to regress the MFE value.

##### Training Data

The UTR-LM model was pre-trained using 5’ UTR sequences sourced from three origins: the Ensembl database, synthetic libraries from Sample et al. [60], and endogenous human 5’ UTR data analyzed by Cao et al. [7].

The preprocessing of 5’ UTR sequences for UTR-LM involved a 4-step pipeline: First, all coding sequence (CDS) and non-5’ UTR fragments were removed from the raw sequences. Second, duplicate sequences were identified and removed. Third, the sequences were truncated to fit within a range of 30 to 1022 base pairs. Finally, incorrect and low-quality sequences were filtered out.

#### A.2.7 BEACON-B

##### Training Data

We filter 523,934 human ncRNA sequences from the total ncRNA in the RNACentral database [68] as pre-training data. BEACON-B and BEACON-B512 use normal BERT-base [21] architecture with 12 layers.

The pre-training configs of BEACON-B and BEACON-B512 are shown as Tab 9 and Tab 10

**Table 9:** Configuration settings for the BEACON-B pre-training.

| config | value |
| --- | --- |
| optimizer | AdamW |
| optimizer epsilon | 1e-6 |
| optimizer momentum | $\beta_1, \beta_2 = 0.9, 0.98$ |
| weight decay | 0.01 |
| learning rate sch. | linear decay |
| learning rate | 2e-4 |
| warmup steps | 10000 |
| steps | 80000 |
| batch size | 256 |
| gradient accumulation | 2 |
| dtype | float16 |
| length | 1026 |
| pertaining data | RNACentral Human ncRNA |

**Table 10:** Configuration settings for the BEACON-B512 pre-training.

| config | value |
| --- | --- |
| optimizer | AdamW |
| optimizer epsilon | 1e-6 |
| optimizer momentum | $\beta_1, \beta_2 = 0.9, 0.98$ |
| weight decay | 0.01 |
| learning rate sch. | linear decay |
| learning rate | 2e-4 |
| warmup steps | 10000 |
| steps | 80000 |
| batch size | 512 |
| gradient accumulation | 1 |
| dtype | float16 |
| length | 512 |
| pertaining data | RNACentral Human ncRNA |
| attention | FlashAttention |

#### A.3 Computational Resources

We fine-tune or train each model from scratch on one task using one NVIDIA A100 40g GPU. We pre-train the simple BEACON-B on 8 A100 GPUs of 80GB for 1.3 days and BEACON-B512 on 4 A100 GPUs of 80GB for 0.895 days.

### A.4 Additional results

For the experiments 4 on component analysis of the baseline RNA language model, we further counted the number of top performances for different tokenizers and positional encoding as shown in Tab 11. The effectiveness of Single nucleotide tokenizer and ALiBi can be demonstrated directly.

**Table 11:**
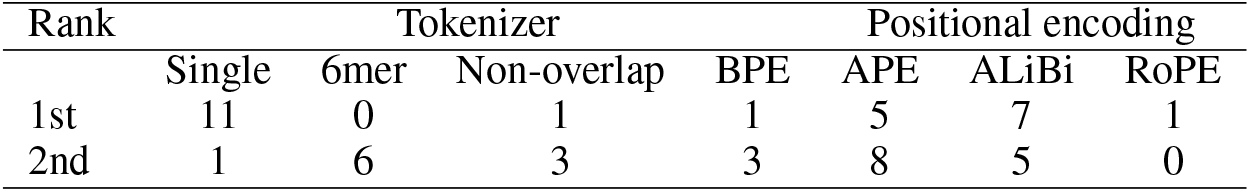
The number of Top2 performance among different tokenizers and positional encodings.

| Rank | Tokenizer |  |  |  | Positional encoding |  |  |
| --- | --- | --- | --- | --- | --- | --- | --- |
|  | Single | 6mer | Non-overlap | BPE | APE | ALiBi | RoPE |
| 1st | 11 | 0 | 1 | 1 | 5 | 7 | 1 |
| 2nd | 1 | 6 | 3 | 3 | 8 | 5 | 0 |

### A.5 Broader Societary Impacts

This work is dedicated to establishing a robust and versatile benchmark for RNA-related tasks, enhancing the understanding of RNA sequences across diverse applications. Our benchmark, encompassing a variety of RNA tasks, aims to rigorously evaluate the efficacy of different RNA representation covering structural analysis, functional studies, and engineering applications. By doing so, it provides a critical assessment of their potential utility in real-world scenarios, thereby laying a foundational framework for applying deep learning in fields such as medical research and genetics.

However, it is also important to acknowledge the dual-use nature of any powerful technology, including those developed from our benchmark. For instance, the enhanced ability to manipulate RNA sequences might be misused, such as in the creation of adverse viral agents. Moving forward, it is crucial to address these risks. We will develop and implement guidelines for the ethical and safe use of our benchmark in the future, ensuring that it contributes positively to society and does not enable harmful applications.

### A.6 Assets

#### A.6.1 Software and Libraries

The open-source software, and corresponding licenses are presented in Tab. 12. The data, licenses and corresponding URL are presented in Tab. 13.

**Table 12:** Software used in this work.

| Asset | License |
| --- | --- |
| FlashAttention [20, 19] | BSD-3-Clause |
| Pytorch [5] | BSD-3-Clause |
| Pytorch Lightning [23] | Apache-2.0 |
| Huggingface [72] | Apache-2.0 |
| Scikit-Learn [49] | BSD-3-Clause |
| Numpy [31] | BSD-3-Clause |
| Matplotlib [34] | Matplotlib License |
| Seaborn [70] | Apache-2.0 |

**Table 13:**
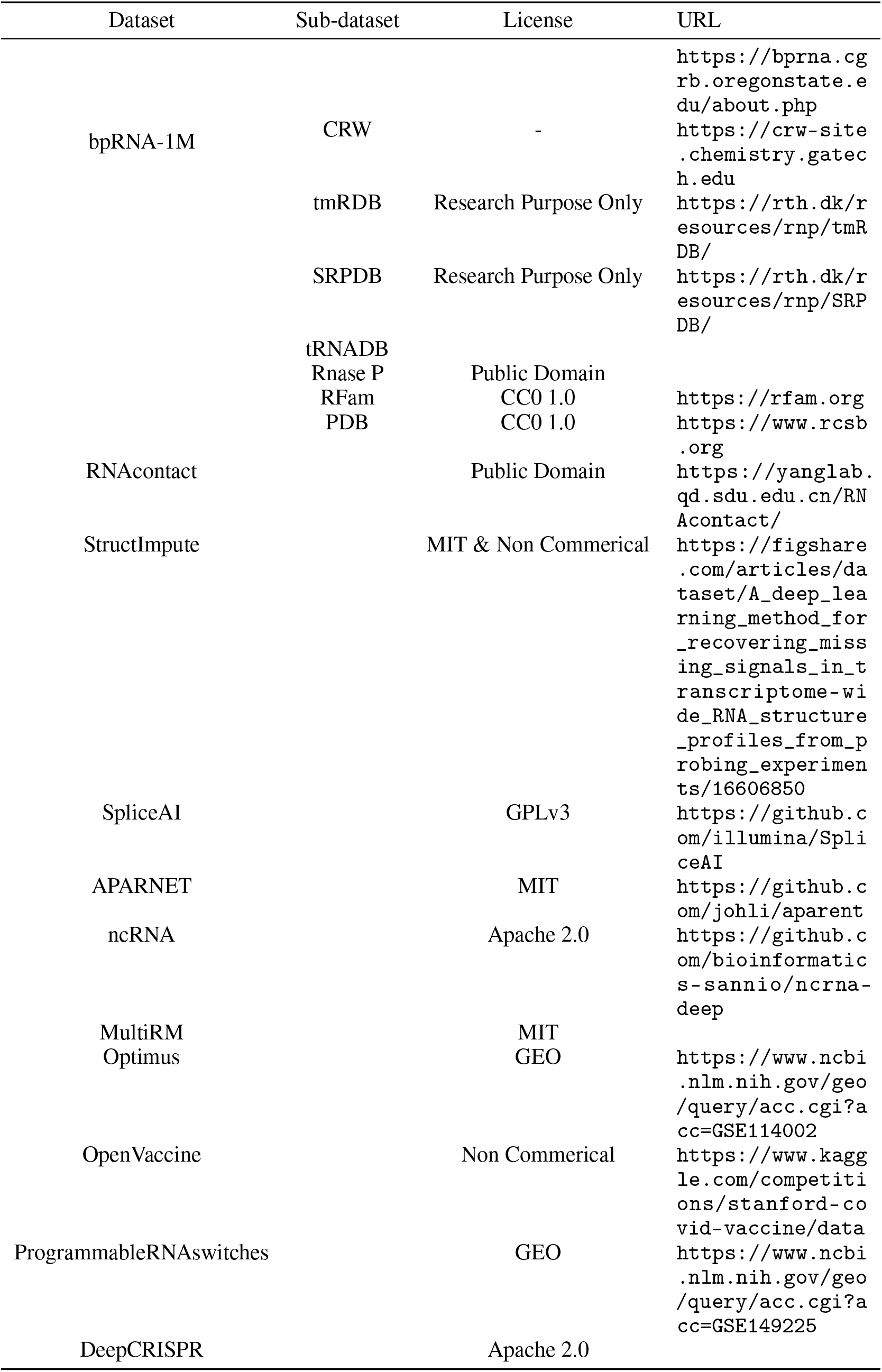
Dataset used in this work.

